# The coupled dual-oscillator model of wing and haltere motion in flies

**DOI:** 10.1101/2020.03.08.982520

**Authors:** Tanvi Deora, Sanjay P. Sane

## Abstract

The mechanics of Dipteran thorax is dictated by a network of exoskeletal linkages which, when deformed by flight muscles, generate coordinated wing movements. In Diptera, forewings power flight, whereas hindwings have evolved into specialized halteres which provide rapid mechanosensory feedback for flight stabilization. Although actuated by independent muscles, wing-haltere motion is precisely phase-coordinated at high frequencies. Because wingbeat frequency is a product of wing-thorax resonance, wear-and-tear of wings or thorax should impair flight ability. Here, we show that wings and halteres are independently-driven, linked, coupled oscillators. We systematically reduced wing length in flies and observed how wing-haltere synchronization was affected. The wing-wing system is a strongly-coupled oscillator, whereas wing-haltere system is weakly-coupled through mechanical linkages which synchronize phase and frequency. Wing-haltere link is unidirectional; altering wingbeat frequency affects haltere frequency, but not vice-versa. Exoskeletal linkages are thus key morphological features of Dipteran thorax, ensuring robust wing-haltere synchrony despite wing damage.

## Introduction

Flies are among the best exemplars of aerial agility. The Dipteran order encompasses a vast repertoire of flight types – from the exquisite hovering and maneuvering ability of hoverflies, to the stable trajectories of mosquitoes and rapid territorial chases in houseflies (Land and Collett, 1974). Such complex maneuvers require precise and rapid control, guided by sensory feedback from multiple modalities (Bender and Dickinson, 2006; Heide and Götz, 1996; Hengstenberg, 1993; Pringle, 1948; Sherman and Dickinson, 2003; Trimarchi and Schneiderman, 1995). Of particular importance for flight stability is the mechanosensory feedback from halteres - the modified hind wings of flies - which sense gyroscopic forces during aerial maneuvers (Nalbach, 1993, 1994; Nalbach and Hengstenberg, 1994; Pringle, 1948). During flight, halteres oscillate in a constant plane at frequencies that are identical to their flapping wings, and with a constant relative phase difference. During an aerial turn, an externally imposed change in the plane of haltere oscillation is resisted due to rotational inertia, causing Coriolis torques to act on the haltere base. Mechanical strain in the haltere shaft due to Coriolis torques is sensed by multiple fields of campaniform sensillae distributed around its base. These encode the stroke-by-stroke status of aerial rotations and provide mechanosensory feedback to the wing muscles (Chan and Dickinson, 1996; Fayyazuddin and Dickinson, 1996; Yarger and Fox, 2018). The relative phase difference between the feedback from wing and haltere mechanosensors determines the activity patterns in wing steering muscles (Fayyazuddin and Dickinson, 1999; Fox et al., 2010). During flight, the two wings of flies move exactly in-phase relative to each other, whereas halteres move at a constant phase relative to wings. This precise phase coordination is maintained at wingbeat frequencies that far exceed 100 Hz (Deora et al., 2015; Hall et al., 2015). Because even slight asymmetries in the bilateral wing motions can result in significant instabilities during flight (Fry et al., 2003), the phase and frequency synchronization of the wing-haltere kinematics is a core feature of Dipteran flight, any deviation from which may signal a self-generated aerial turn or an unwanted perturbation.

Previous research has shed much light on the architecture of the Dipteran thorax (Deora et al., 2017; Ennos, 1987; Miyan and Ewing, 1985; Pringle, 1949). The wing-haltere system acts as a complex resonant box. In flies, wings are actuated by two sets of antagonistic indirect flight muscles aligned dorso-longitudinally and dorso-ventrally within the thorax (Deora et al., 2017; Dickinson and Tu, 1997; Pringle, 1949). Their activation is myogenic; hence, contraction in one set of muscles triggers delayed contraction of the other and vice-versa, setting up resonant cycles of oscillations of the entire thorax. A complex wing hinge transforms oscillatory deformations of the thorax into large-amplitude wing strokes. Indirect flight muscles require neural stimulation to remain in an active state, but the frequency of stimulation is typically an order of magnitude lower than resonant thoracic oscillations. The attitude of the wing is finely adjusted on a stroke-to-stroke basis by a set of steering muscles which are under direct neuronal control (Lindsay et al., 2017). Thus, frequency characteristics of Dipteran wing movements are set by the resonance frequency, which in turn is determined primarily by the wing-thoracic morphology. Unlike wings, the motion of each haltere is powered by a single asynchronous muscle whose contractions activate its upstroke, whereas the downstroke is thought to be entirely passive (Pringle, 1949).

Although wingbeat frequency depends on wing-thorax morphology, the precise coordination of wings and halteres is mediated by mechanical linkages within the thorax which ensure tight coupling of phase and frequency (Deora et al., 2015). This suggests the hypothesis that wings and halteres act as independent forced oscillators, whose kinematics are both coupled and constrained by two separate mechanical linkages within the thorax (*the coupled dual-oscillator hypothesis*, figure 1A). One linkage, *the scutellar link* (alternatively, *wing-wing link*), is embedded within the scutellum and ensures that both wings are synchronized. A second linkage, *the sub-epimeral ridge* (alternatively, *wing-haltere link*), ensures precise coordination of wings and halteres. This linkage ensures only weak-coupling of wing-haltere synchrony, but breaks down when wingbeat frequency exceeds a threshold value (Deora et al., 2015).

**Figure 1:**
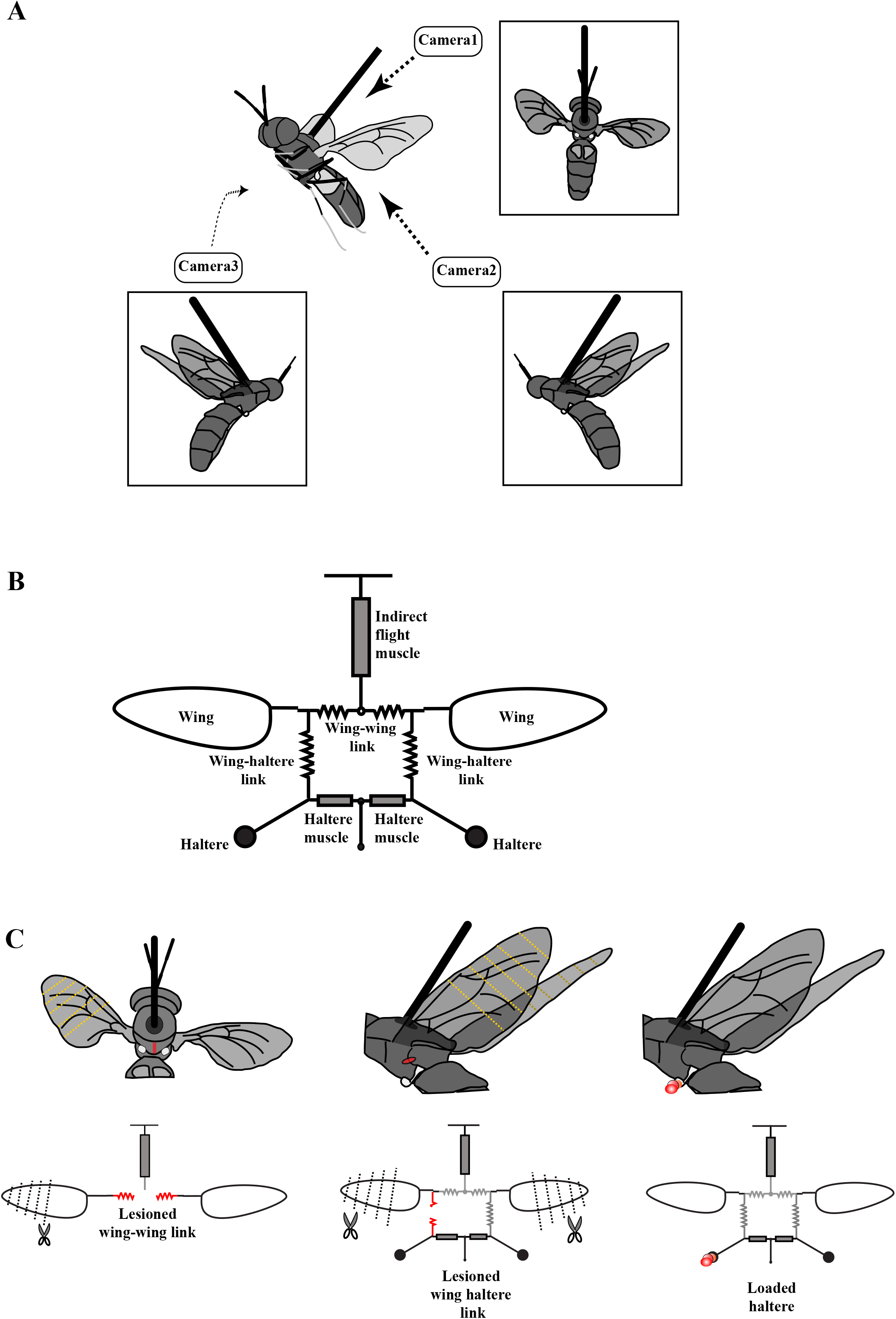
The experimental setup. (A) Schematic of a tethered fly showing the position of the three high-speed cameras. Insets show the three different camera views. (B) Mechanical model of the Dipteran thorax modified from an earlier work (Deora et al., 2015). This model excludes the clutch and gearbox from the previous figure, focusing instead on the wing-wing and wing-haltere linkages which are the main focus of the experiments described here. (C) A schematic (top) and model (bottom) illustrating the experimental treatments. The treatments (red) included lesioning the scutellum or wing-wing link (left panel), lesioning sub-epimeral ridge or wing-haltere link (middle panel) and haltere-loading (right panel). The clipped wings are also indicated (dotted lines). The same treatments are also shown in the model schematic, and used as insets in later figures.

The resonant properties of such a system rely on mechanical integrity of the wing-thorax system. However, wings of insects often undergo significant natural wear-and-tear during the lifetime of an adult insect (Hayes and Wall, 1999). Wing damage alters both frequency and aerodynamic force generation of the flapping wings (Hedenström et al., 2001), thus posing a challenge to the overall coordination of wing motion. Such damage is typically asymmetric and can affect maneuverability. Not surprisingly, in insects such as bumblebees and dragonflies, wing damage leads to decreased success in hunting and also greater mortality (Cartar, 1992; Combes et al., 2010; Haas and Cartar, 2008).

Here, we address two related questions. First, how do flies maintain symmetric wing movement under conditions of wing damage? Second, if wings and haltere motion follows the coupled dual-oscillator hypothesis, how robust are the wing and haltere kinematics in face of wing or thoracic damage? To address these questions, we conducted a series of experiments on the soldier fly, *Hermetia illucens*, in which we made specific lesions of scutellar linkages, and sub-epimeral ridge to impair the mechanical integrity of the thorax. In addition, we either clipped the wings or loaded the halteres to alter their oscillation frequencies. These experiments enabled us to systematically test the predictions of the coupled dual-oscillator hypothesis, and outline the key mechanical properties of the Dipteran thorax that ensure wing-haltere coordination.

## Methods

### i) Surgical treatments and tethering procedure

1-4 days old soldier flies were cold-anesthetized by placing them in an ice box for 5 minutes. We performed surgical treatments (see figure 1B) before their recovery from cold anesthesia:

a. In *Control flies*, no surgeries were performed but they were otherwise handled in the same way as experimental flies.
b. In *scutellum-lesioned flies*, we made a small cut only in the scutellum using a scalpel blade (#11, Fine Science Tools Inc., Foster City, CA, USA) while leaving rest of the thorax intact (left panel, figure 1B).
c. In *unilateral sub-epimeral ridge lesioned flies*, we lesioned the sub-epimeral ridge at a position anterior to the spiracle on the left side of thorax (middle panel, figure 1B). The right ridge was kept intact and served as internal control. To examine how sub-epimeral ridge influences wingbeat frequency, we compared data from *unilateral sub-epimeral lesioned* group with previously published data on *control* flies (Deora et al., 2015). In both cases, procedures for rearing, handling and tethering were identical.
d. In *Bilateral sub-epimeral ridge lesioned flies*, we lesioned sub-epimeral ridges on both sides.
e. In *Bilateral haltere ablated flies*, we cut out the knob of halteres on both sides.

Post-surgery, we tethered the insects with cyanoacrylate adhesive to the tip of a needle bent to 1~90° while they were kept on a pre-chilled metal block. The bent tip provided the necessary surface area to glue a fly to the tether. The tether was lowered using a 3-way micro-manipulator (Narishige Scientific Instrument Laboratory, Tokyo, Japan) and attached dorsally on the scutum of the fly. Flies were given at least an hour for recovery before recording their flight.

### ii) Wing-haltere perturbations and filming procedure

We positioned flies at about 60° to the horizontal (approximately its position during free-flight), and elicited flight by lightly touching their abdomen with a brush. We used three high-speed cameras (v7.3 Phantom camera, Vision Research Inc, Wayne, NJ, USA) to film the insects in flight at a resolution of 800×600 pixels at 2000 frames per second (approximately 15-20 times the wingbeat frequency) and 100 μsec exposure. The three cameras (one top view and two side views) captured the 3D motion of both pairs of wings and halteres (figure 1C). The three camera views were calibrated for each filming bout using a custom-made calibration object.

#### Flies with reduced wing length

We first filmed the flight of flies with intact wings, and then switched off the lights to inhibit their flight. Under a dissection microscope attached with a light source and a red-filter to cut off wavelengths below 610 nm, we clipped their wings with a pair of scissors (Fine Science Tools Inc, Foster City, CA, USA) to an appropriate length using wing vein patterns as landmarks. To test the coupled dual-oscillator model (figure 1A), we clipped one wing (either left or right) while keeping the other intact (figure 1B). In all other experiments, both wings were symmetrically reduced. After each round of wing clipping, we filmed the experimental fly in flight. Each series of experiments yielded 4-6 data points including 1 intact and 3-5 reduced wing lengths.

#### Flies with loaded halteres

We initially filmed flies with intact wings and halteres, which served as control. Next, we loaded the left haltere with varying amounts of glue and metal shavings under a dissection microscope under red light. Because of the small size of halteres, the amount of load could not be accurately quantified, but we systematically decreased the haltere frequency by incrementing an arbitrary amount of glue and load mixture (right panel, figure 1B). First, we loaded the haltere with glue (Fevicol™; polyvinyl acetate, Pidilite Industries) mixed with red poster paint using a metal insect pin. We next added increasing amounts of aluminum shavings mixed with glue and paint to the already loaded haltere. Following each load increment, we filmed the tethered fly. After three rounds of loading, we carefully removed the film of glue, paint and metal shaving with a pair of forceps under the microscope and again filmed the flight recovery. In a few cases, flies removed the load while self-grooming. This experimental procedure yielded 5 data points for each fly – 1 intact, 3 increasing amounts of load and 1 with the load removed.

## Results

### Asymmetric wing damage does not alter wing coordination

The left and right wings are coupled by a mechanical linkage running through the scutellum within the thorax such that the two wings always flap at constant phase relative to each other (figure 1A) (Deora et al., 2015). However, mechanical coupling of the two wings also implies that their flapping frequencies are identical, and thus alteration in the frequency of one wing should correspondingly alter frequency of the contralateral wing. To test this prediction, we filmed the wing motion of tethered soldier flies in which the length of one wing was sequentially reduced while leaving the other intact. Wingbeat frequency of one-wing clipped fly was determined by sampling 20 arbitrarily chosen wing beats. Even after drastic reduction in the length of one wing by >50% of the original length, the frequency of the clipped and intact wings was always identical (figure 2A left and 2B; for additional data see figure S1, p=0.623; Wilcoxon signed-rank test on the maximum wing difference). Thus, the overall wingbeat frequency is determined by the frequency of the intact wing.

**Figure 2:**
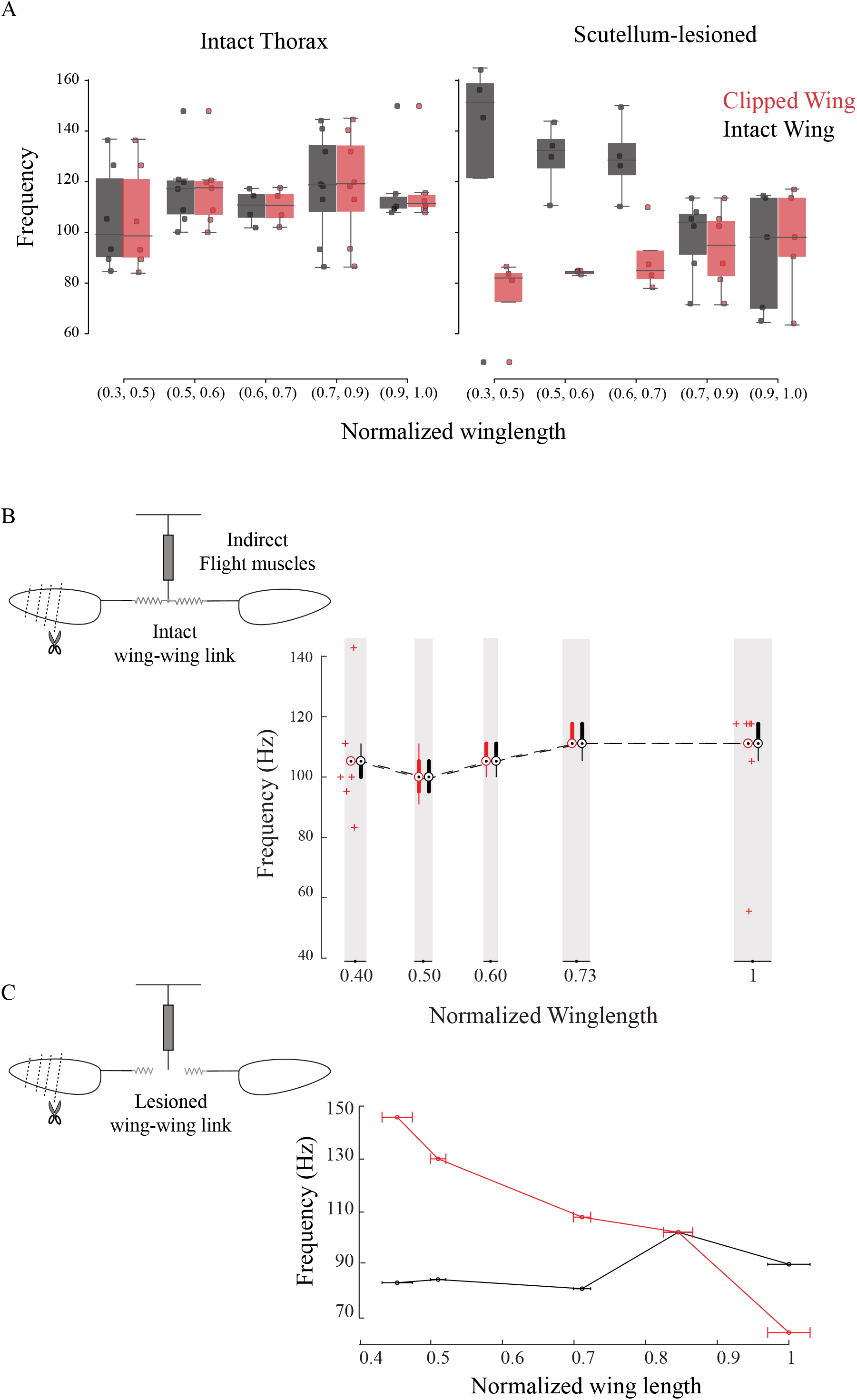
The frequency of the two wings is synchronized by the scutellum. (A) Wingbeat frequency of intact (grey) and clipped (red) wing as a function of clipped-wing length for intact thorax (left panel) and slit scutellum (right panel). Flies with intact thorax flap their wings at identical frequencies whereas the *scutellum-lesioned* flies’ flap at different frequencies. Each dot represents an individual fly. (B) Wingbeat frequency as a function of clipped wing length for a representative, intact fly plotted as compact box plot (~20 wingbeats are analyzed at each wing length). (C) Peak wingbeat frequency as a function of clipped wing length for a representative *scutellum-lesioned* fly. Insets show schematic for treatments. Additional data for individual flies can also be found in SF 1 and 2.

### Scutellar integrity is essential for wing coordination

According to the coupled dual-oscillator model, lesioning the scutellum should decouple the frequencies of left and right wings. Accordingly, we severed the scutellar linkage and filmed the flies while again sequentially clipping one wing to reduce its aerodynamic resistance thereby increasing its frequency. This resulted in irregular wing beats in these flies, with frequent mid-stroke pauses and an overall reduction in stroke amplitude. Fourier analysis of the time series of wing motion shows that both wings oscillated at very different frequencies. Thus, unlike the intact scutellum case in which frequency synchronization was robust to wing damage (Fig 2A left panel and Fig 2B), the wings of a scutellum-lesioned fly were decoupled from each other (figure 2A right panel and 2C, additional data in figure S2, p=0.022; Wilcoxon signed-rank test on the maximum wing difference). For example, in the typical case of a scutellum-lesioned fly, the clipped wing flapped at 130 Hz when cut to 50% of its original length, as compared to 85 Hz in the intact wing (figure 2C). Lesioning the scutellum disrupts, and reattaching completely restores the phase coordination between both wings (Deora et al., 2015); thus, scutellar integrity is essential to ensure precise coupling of both phase and frequency of the two wings, and imparts robust synchronization even if one or both wings are slightly damaged or torn. These data show that the two wings are strongly coupled by the scutellar link.

### Sub-epimeral ridge weakly couples the frequency of each haltere to its ipsilateral wing

The wing and haltere motion on each side is coupled by a separate thoracic element called the sub-epimeral ridge (figure 1B). A small reduction in wing length resulted in a small increase in wingbeat frequency, and concomitant increase in haltere frequency. However, with further reduction of wing length, wingbeat frequencies exceeded ~150% of the initial values, and halteres failed to keep pace with the wings. In such conditions, haltere frequency dropped closer to their natural frequency suggesting that their coupling was weak (figure 3, control haltere (in blue)) (Deora et al., 2015).

**Figure 3:**
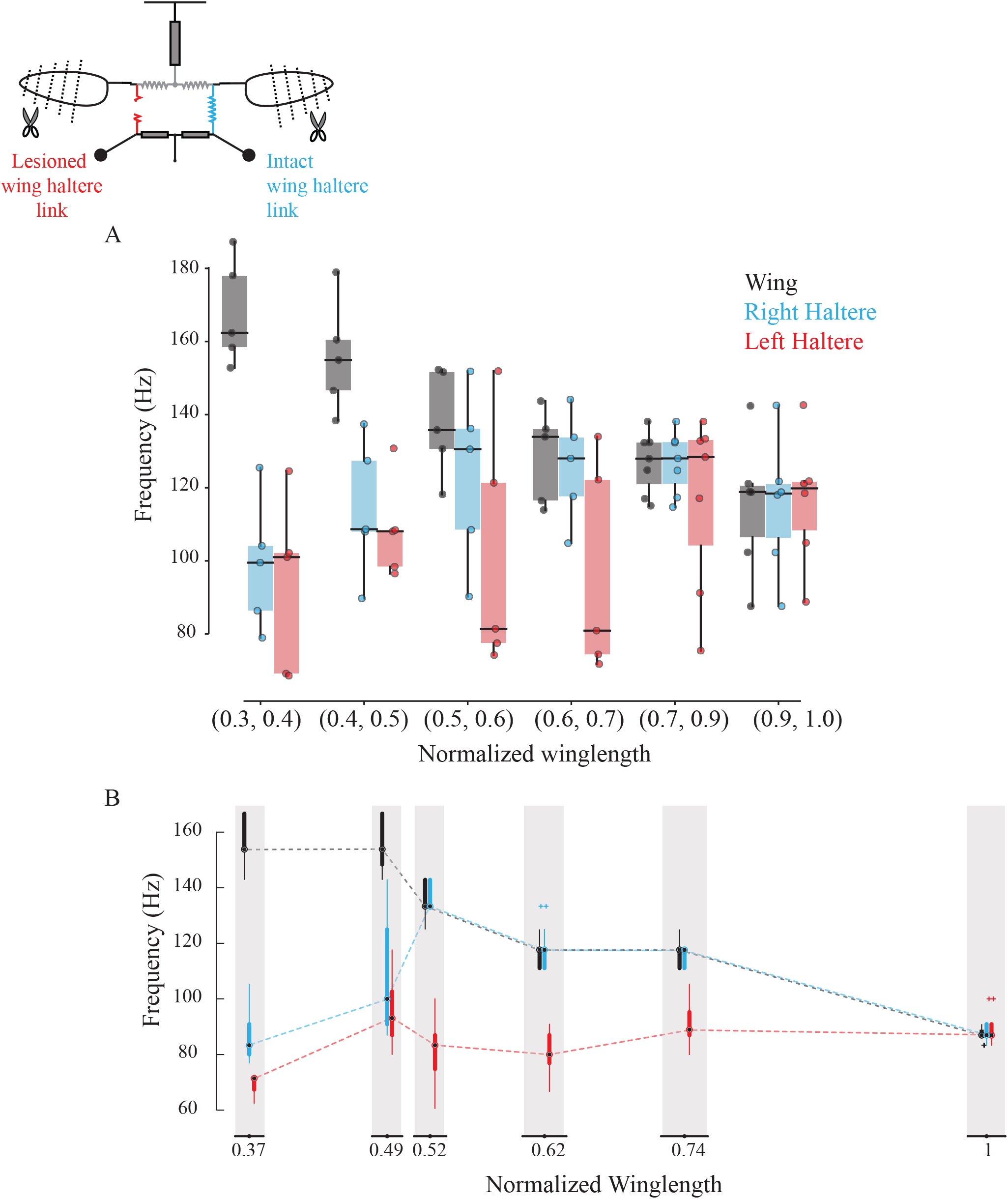
Sub-epimeral ridge couples the frequency of wings and halteres. Frequency of wing (grey), control haltere (blue) and haltere with the sub-epimeral ridge lesioned (red) as a function of wing length across all flies (A) and one representative fly (B). Inset shows the schematic for treatments. Additional data for individual flies can be found in SF3.

We next lesioned the sub-epimeral ridge on the left side while keeping the right side intact as internal control. If sub-epimeral ridge is the main coupling link, then lesioning it should cause the haltere frequency on the lesioned (left) side to be decoupled from the increase in wingbeat frequency due to wing shortening. Our data were consistent with this hypothesis; the control (right) haltere frequency matched the wing frequency more robustly than the lesioned left haltere-wing pair (Fig 3A, p=0.039 for wing-treatment haltere pair and p = 0.657 for wing-control haltere pair, one-sided Wilcoxon signed-rank test at wing length bin = [0.6, 0.7]). Not surprisingly, these data were more variable. In 4 out of the 6 experiments, data were consistent with our hypothesis; the frequency of the haltere on the lesioned (left) side either did not increase at all (representative fly in figure 3B, figure S3 A & B) or was decoupled from the wing even with slight changes in wing length, thus displaying no robustness in the wing-haltere synchrony (figure S3C). In 2 flies, however, wingbeat frequency remained relatively unchanged despite clipping the wings incrementally, and haltere frequency on the lesioned side matched halteres on the control side (figure S3 D & E). Together, these results suggest that the sub-epimeral ridge weakly couples wing and haltere oscillation. Haltere motion can accommodate small to moderate changes in wingbeat frequency but fails if these changes are large.

### Integrity of the sub-epimeral ridge is essential for resonant oscillation of the thorax

In insects with an intact thorax, clipping the wings increases wingbeat frequency by as much as 90 Hz. In flies with unilaterally-lesioned sub-epimeral ridge, changes in wingbeat frequency were relatively moderate (~60 Hz) after wing shortening (figure 4, data for intact flies from Deora et al., 2015, p<0.05, Kruskal-Wallis ANOVA followed by the post hoc Tukey-Kramer multi-comparison test, see supplemental methods for additional details).

**Figure 4:**
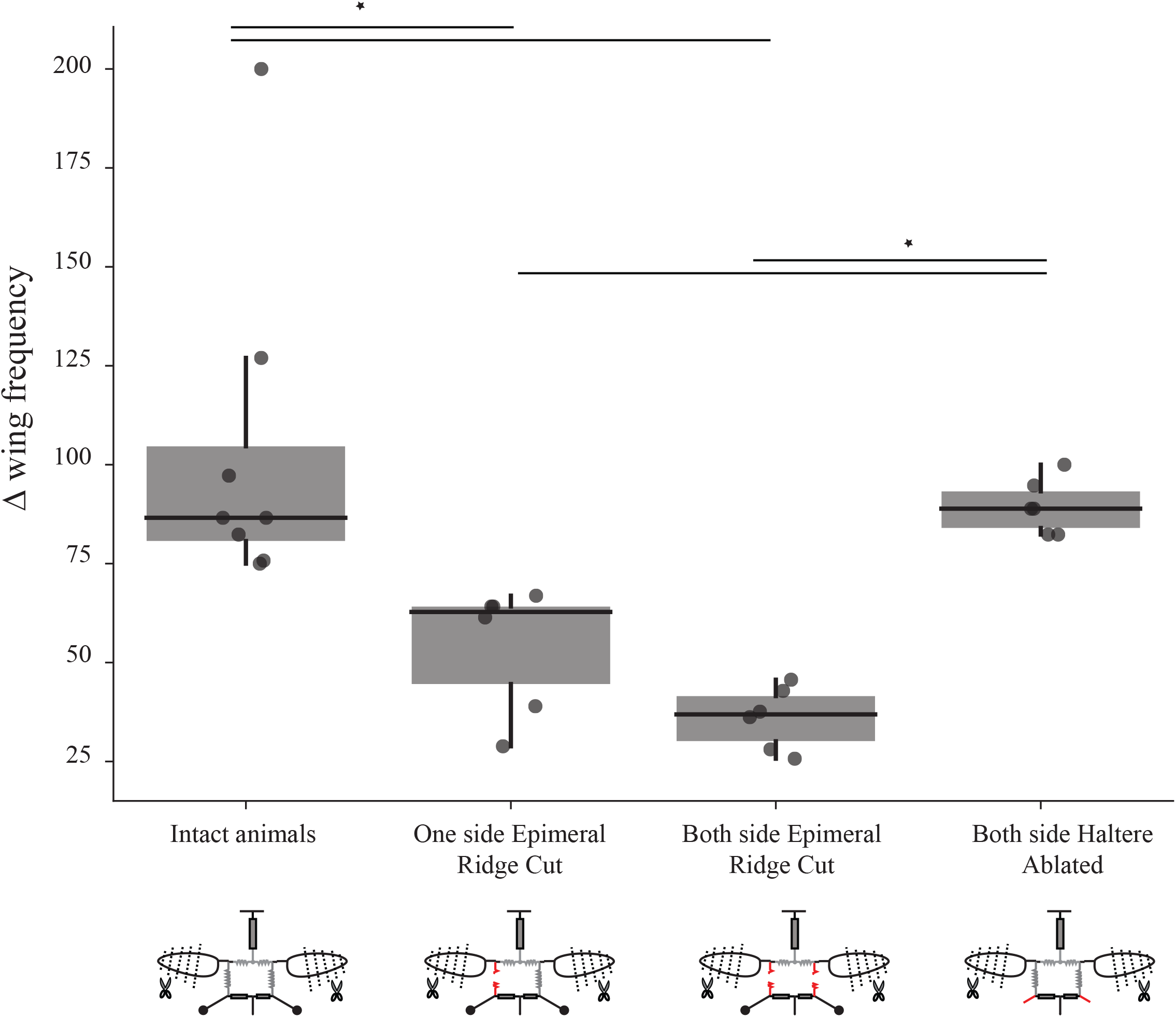
Sub-epimeral ridge lesions alter the resonant properties of the thorax. Individual box plots show increase in wing beat frequency for reduced wing length in different treatment groups. Flies with one or both sub-epimeral ridge lesioned show a smaller increase in wing beat frequency as compared to intact flies. In contrast, flies with both haltere ablated can increase their wing beat frequency significantly more than flies with both sub-epimeral ridges lesioned. (* p value < 0.05, non-parametric Kruskal-Wallis ANOVA followed by a Tukey-Kramer post hoc multi-comparison test, n; 8 intact flies, 6 flies for the other three treatments). Insets under each group show the schematic for treatments.

How does lesioning the sub-epimeral ridge alter wingbeat frequency, in addition to decoupling the wings and halteres? One possibility is that a lesioned ridge disrupts the anti-phase motion of wings and halteres, leading to aberrant haltere feedback to wing steering muscles thereby affecting wingbeat frequency. Alternatively, a lesioned ridge could mechanically disrupt frequencies by acting as a free end that dissipates energy, thereby disrupting the overall resonant mechanics of the thorax.

To test for these possibilities, we first lesioned the ridges on both sides. In these flies, the increase in wingbeat frequency was even further restricted (<40 Hz). In another set of flies, we kept both the sub-epimeral ridges intact but ablated both halteres. Ablating halteres alters their feedback but does not mechanically disrupt the thoracic linkage network. Clipping wings of the haltere-ablated flies resulted in significantly elevated wingbeat frequency (by ~90 Hz, figure 4) as compared to flies with both sub-epimeral ridges lesioned (p <0.05) but similar to flies with an intact sub-epimeral ridge. This suggests that mechanical integrity of these linkages most likely determines wingbeat frequency. Hence, the mechanical integrity of the entire thoracic linkage system, including both the scutellum and sub-epimeral ridge is essential to maintain the resonant properties of the thorax.

### Sub-epimeral ridge is a unidirectional linkage

If the sub-epimeral ridge is bidirectional, then wing frequency should change when haltere frequency is experimentally altered, and vice-versa. Halteres, like wings, are powered by myogenic musculature, and thus changing the haltere mass affects its frequency. To test the hypothesis that sub-epimeral linkage is bidirectional, we loaded halteres with small weights thereby altering their frequency, and measured the effect on wingbeat frequency. Unlike the wing clipping, we could not reduce the haltere mass in discrete steps as it is mostly concentrated at the end knob. Instead, we sequentially loaded each haltere knob with small amounts of glue, thus reducing its frequency in discrete steps (Methods (ii); figure S4 A-C).

Small loads did not affect haltere frequency, but as the load increased beyond a threshold, haltere frequency decreased in discrete steps. However, wingbeat frequency remained constant in these experiments (figure 5; for individual fly data see figure SF5A-F, p = 0.014; Wilcoxon signed-rank test for left loaded haltere and wing pair). Thus, the sub-epimeral ridge is a unidirectional link, which couples haltere motion to wing motion but not vice-versa. To rule out the possibility that loading the haltere irreversibly damaged haltere muscles or the sub-epimeral ridge, we detached the load and confirmed that haltere frequency recovered and again matched wingbeat frequency (figure 5; S5).

**Figure 5:**
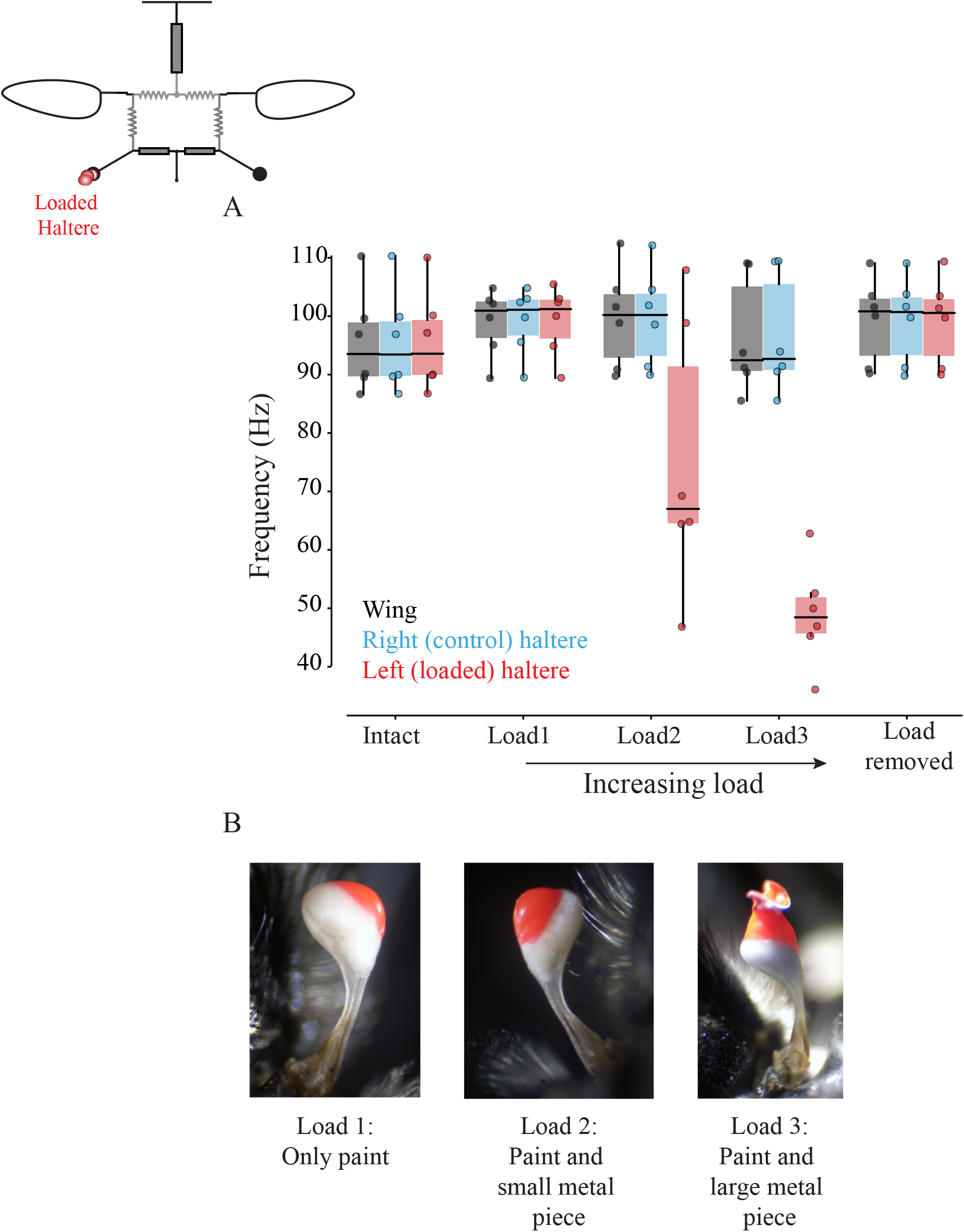
Wing-haltere linkage is unidirectional because changing haltere frequency does not alter wingbeat frequency. Frequency of wing (grey), control haltere (blue) and loaded haltere (red) across all flies. Each dot represents a single individual fly. The haltere frequency drops as the haltere is loaded but the wing frequency remains the same showing that wing-haltere coupling is unidirectional. Inset shows the schematic for the treatment. Data for individual flies can be found in SF5. (B) Representative images of the haltere loaded with different amounts of load (also see figure S4).

## Discussion

For stable flight, bilateral wing motion must be precisely coordinated, and halteres must maintain a precise phase difference relative to wings to ensure correct feedback to wing steering muscles (Deora et al., 2015; Fayyazuddin and Dickinson, 1999; Fry et al., 2003). Here, we show that the phase and frequency of wings and haltere motion is mechanically coupled by thoracic linkages, thereby imparting robustness of wing-wing and wing-haltere coordination against damage or wear-and-tear.

### Mechanical linkages enable robust frequency-phase output despite asymmetric wing damage

Physical damage to the wings of adult insects is irreversible, and potentially deleterious for fitness. Wings of certain insects have flexible costal break or specific venation patterns that prevent wing damage (Mountcastle and Combes, 2014). However, despite such adaptations, insects incur wing damage in the wild due to factors like predation and age (Cartar, 1992; Hayes and Wall, 1999) When one or both wings are damaged, the overall aerodynamic load reduces thereby increasing the frequency of thoracic and wing oscillations (Deora et al., 2015). The linkages described here ensure that the phase and frequency matching between wings and halteres is robust despite significant changes in wingbeat frequency, and hence they may be viewed as evolutionary adaptations that impart robustness to wing motion in case of damage or wear-and-tear.

Natural wing damage is typically asymmetric. Shortening one wing by as much as 50% did not significantly alter the resonant frequency of thorax, perhaps because stretch-activation is dictated primarily by the wing with greater aerodynamic load. Importantly, in flies with an intact scutellar linkage, asymmetric changes in wing length did not alter the overall synchrony; wings remained bilaterally coordinated, regardless of their length (figure 2).

In bees with symmetric or asymmetric wing damage, wingbeat frequency increases during free flight (Hedenström et al., 2001; Vance and Roberts, 2014), perhaps due to reduced wing inertia. Both bees and flies have similar indirect, asymmetric flight muscle architecture. However, contrary to freely flying bees, in our tethered experiments asymmetric wing damage in flies with intact thorax did not increase wingbeat frequency, indicating that insects actively maintain their wingbeat frequencies. However, lesioning the scutellum completely decoupled the frequencies of the two wings of different lengths; shortened wings oscillated at frequencies up to twice that of the intact wing. These results underscore the importance of the scutellar linkage in ensuring precise coordination between the two wings.

### Control of bilateral kinematics by indirect flight muscles

When one wing was shortened and the scutellar link was lesioned, the shortened wing flapped at higher frequency than the intact wing. This shows that the indirect flight muscles on the two sides are, in principle, capable of operating at different frequencies, but are constrained to flap synchronously by the scutellar link. These results are consistent with the idea that indirect flight muscles aid the direct flight muscles in power modulation and kinematic control of wings (Gordon and Dickinson, 2006; Lehmann et al., 2013). In intact flies, power modulation occurs under constraints of equal bilateral wingbeat frequency and phase which leaves open the possibility that amplitude or stroke plane can be modulated by power muscles. For instance, during turns, the two sets of dorso-longitudinal muscles can be differentially activated by their motor neurons (Gordon and Dickinson, 2006; Lehmann et al., 2013), potentially leading to differential power output. Moreover, wing damage results in differential force production on the contralateral sides in hawkmoths (Fernández et al., 2012). Asymmetric wing damage in hawkmoths causes activation delay in the dorso-ventral power muscle, resulting in a voluntary yaw-like maneuver towards the undamaged side, perhaps to compensate for the reduced lift on the damaged side. However, it is important to note that although hawkmoths have indirect flight muscles, they are synchronous and hence under direct neural control. Our results suggest that, like hawkmoths, the indirect, asynchronous flight muscles of flies could also modulate power output of the ipsilateral wing, independent of the contralateral power muscles.

### Role of sub-epimeral ridge in wing-haltere coordination

The sub-epimeral ridge synchronizes the frequency and phase difference between wings and halteres on each side (figure 3, (Deora et al., 2015)). How would the structural diversity of the thorax and linkages in diverse Diptera influence wing-haltere motion? For example, as compared to the rounded thorax with an almost circular sub-epimeral ridge in blowflies, the thorax of mosquitoes is thinner with an oblong sub-epimeral ridge. Presumably, there is also variation in the material properties and strain transfer across the cuticle. Across flies, there are also differences in their haltere kinematics. For example, blowfly halteres flap exactly antiphase to the wings whereas, for mosquitoes this phase is closer to 0.

It is not clear how the precise phase difference is set in these flies, but these parameters are likely the outcome of the variation in the thoracic and linkage anatomy across Diptera and the physics of coupled, driven oscillator system. Indeed, when wing and haltere frequencies are decoupled by loading the haltere (figure 5), the haltere oscillates at an altered phase relative to the wing, even when sub-epimeral ridge is intact (figure S6 & 7). Their frequency, on the other hand, is thought to be determined primarily by the stretch-activation properties of their main driving muscles – the indirect flight muscles for wings, and haltere muscles (or Pringle’s muscle) for halteres. Because the sub-epimeral ridge couples haltere frequency to wingbeat frequency, their vibration frequency is fine-tuned by this linkage and the overall thoracic geometry.

### Weak coupling properties of the sub-epimeral ridge

The sub-epimeral ridge weakly couples wings and halteres. Its stiffness is limited by its material strength and geometry and it can ensure coordination of wing-haltere frequency close to original frequency. At wingbeat frequencies greater than about 150% of the original frequency, the haltere frequency reverts to a value that is closer or equal to its natural frequency. This behavior is typical of independently driven oscillators, coupled by a mechanical element of finite coupling strength (Strogatz, 1994). Moreover, the sub-epimeral ridge acts in a unidirectional manner (figure 5); moderate changes in wingbeat frequency alter the haltere frequency, but not vice-versa. This may be the outcome of the large difference in the wing and haltere masses (and therefore inertia), and their respective muscles. Our experiments show that the aerodynamic load on the wings determines flapping frequency, and the halteres merely follow.

### General implications for other insects with asynchronous muscles

Although our study was focused on Dipteran insects, several implications of this study extend beyond Diptera. Indeed, all insects that possess asynchronous flight muscles must rely on linkages for wing-wing coordination. During buzz pollination in bees and pre-flight thoracic vibrations in beetles the wings are folded even as the flight muscles are active and vibrate the thorax (Esch and Franz, 1991; Leston et al., 1965). Such behaviors suggest the presence of passive mechanical linkages in other insect groups that have asynchronous flight muscles. It is also likely that in many cases, relative coordination between the front and the hind wings are mediated by passive linkages analogous to the sub-epimeral ridge in Diptera. Thoracic linkages may also play a very important role in miniature insects, in which there are fewer muscles groups for higher-level control. Extreme miniaturization in insects of the order Diptera, Hymenoptera and Coleoptera (Polilov, 2015) poses severe physiological and biomechanical constraints on the organism (Polilov, 2012; Sane, 2016). Because smaller wings generate reduced aerodynamic lift, miniature insects must flap at increased frequencies to generate enough lift to stay aloft. In such insects, thoracic linkages are likely to play a key role in mediating synchronization of the wings, while also constraining their degrees of freedom. Small body sizes are associated with rapid wingbeat frequencies, often exceeding 100Hz (Dudley, 2000). The key results in this paper show that the thoracic morphology of the fly plays an important role in providing robust wing coordination despite wing damage. How thoracic morphology and material characteristics are adapted for such rapid, coordinated wing movements remains a fascinating question for future studies.

## Acknowledgments

We would like to thank Prof. Sandeep Krishna, Prof. Rudra Pratap, Rizwana Parween and Akash Vardhan for the discussions on mechanical oscillators and Dr. Anand Krishnan for discussions and help with writing the manuscript.

## Author contributions

SPS and TD designed the experiments; TD performed experiments and analyzed the data; SPS and TD wrote the paper.

## Supplementary Material

### Methods

#### i) Soldier fly rearing procedure

Wild caught black soldier flies, *Hermetia illucens* were enclosed in vials filled with a medium of corn flour and agar mixed with yeast powder. In most cases, we caught wild gravid females which laid eggs immediately upon capture. We reared the larvae on the artificial medium, frequently replacing it as the larvae grew. Through most of the year, adult flies emerged approximately a month after pupation, but their emergence during winters could be more delayed due to diapause. Adult soldier flies were maintained in mesh cages on a 12:12 hour day-night cycle. Some flies were reared on compost in which wild gravid females laid eggs. The larvae fed and pupated in the compost. Pupae were separated and collected in a separate box. Adults were maintained in natural day-night cycle. Animals reared in this manner were typically bigger and more active than lab-reared animals but showed no difference in behavior.

#### ii) Analysis

We computed the time period (and frequency) of a single wing and haltere stroke from videos by counting the number of frames per wing stroke. We analyzed 20 strokes per flight bout at each wing length. In the *scutellum-lesioned group*, the flies often flapped their wings with varying amplitude and frequency making it difficult to ascertain the precise time duration for each wing stroke. We digitized the videos of these animals using the DLTdv3 code (Hedrick, [1]) in MATLAB (Mathworks Inc, Natick, MA, USA) and analyzed the data using custom MATLAB codes. For each wing, we calculated the azimuthal (theta) angle and plotted the theta position of both wings as a function of time and obtained the average wing beat frequency using Fourier analysis.

We calculated the change in wingbeat frequency after clipping wing length (ΔF), as

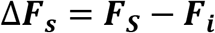

where,

F_s_ = wing beat frequency at the shortest wing length
F_i_ = wing beat frequency at the intact wing length

Increment in wingbeat frequency due to reduced wing length (ΔF) for all four groups was compared using non-parametric Kruskal-Wallis ANOVA followed by the Tukey-Kramer post hoc multi comparison analysis. Codes used for analysis can be found at https://github.com/TanviDeora/Coupled_Dual-oscillator_DeoraSane.

**Table 1:**
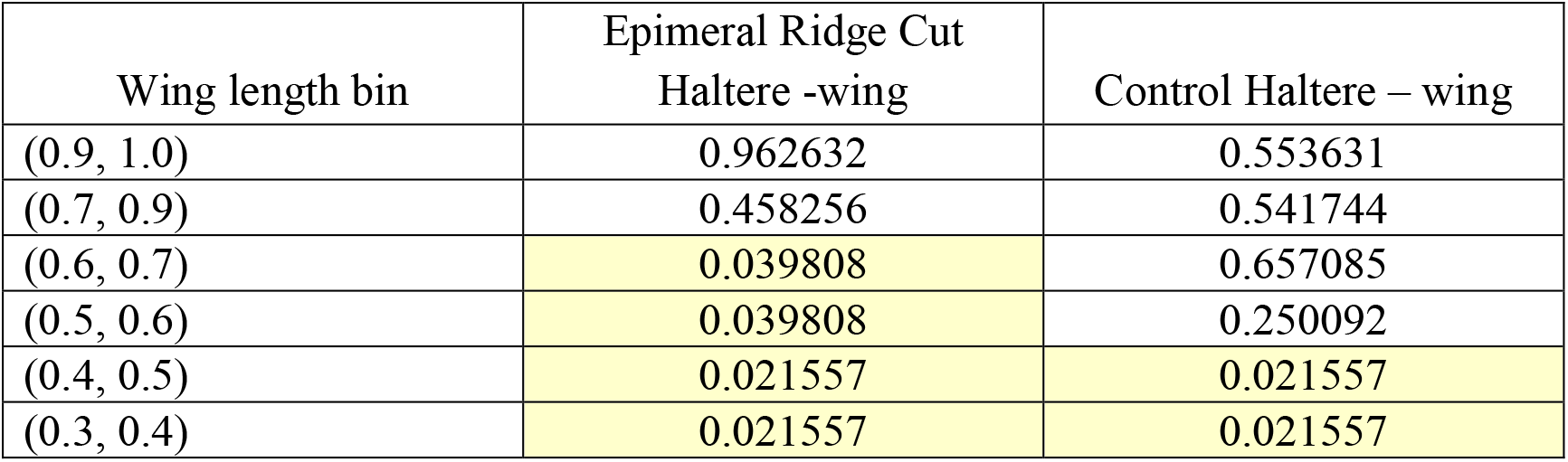
p values for a one-sided Wilcoxon signed rank sum test at all wing length bins for epimeral ridge cut flies in Figure 3A.

**Table 2:** p values for a one-sided Wilcoxon signed rank sum test between haltere and wing pairs at all haltere loads in Figure 5.

|  | Loaded Haltere - wing | Control Haltere - wing |
| --- | --- | --- |
| Intact haltere | 0.875568 | 0.827384 |
| Load 1 | 0.963031 | 0.986146 |
| Load 2 | 0.013854 | 0.662929 |
| Load 3 | 0.013854 | 0.986146 |
| Load removed | 0.231536 | 0.342915 |

### Testing Significance and estimating sample sizes

To estimate a minimum sample size to detect a difference between two groups, we used the estimated mean and std of these two groups (measured from data) and simulated datasets of different sample sizes. For each simulated dataset, we calculated the p value using the relevant test (Wilcoxon signed-rank test for all paired data and Kruskalwallis for groups). We used a bootstrapping method; repeating this about 10,000 times for each sample size and estimated the power, i.e. probability of detecting a difference between these two groups at significance level of 0.05. Below we report the tested groups and power analysis for each experiment.

### Wing Coordination by Scutellum (Figure 2)

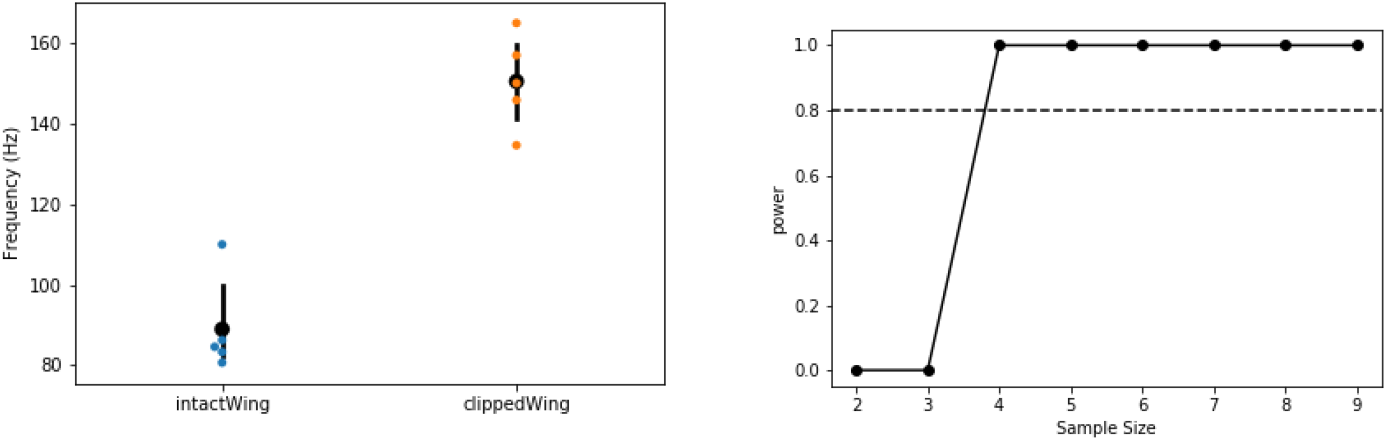

(*Left*) The frequency of intact and clipped wing (at the wing length which has the largest difference) are significantly different (Wilcoxon signed-rank test, p = 0.021). (*Right*) Power analysis for different sample sizes. Our sample size of 6 is greater than the minimum sample size (=4) needed to have 80% confidence level (dashed line)

### Wing-Haltere Coordination by Epimeral Ridge (Figure 3)

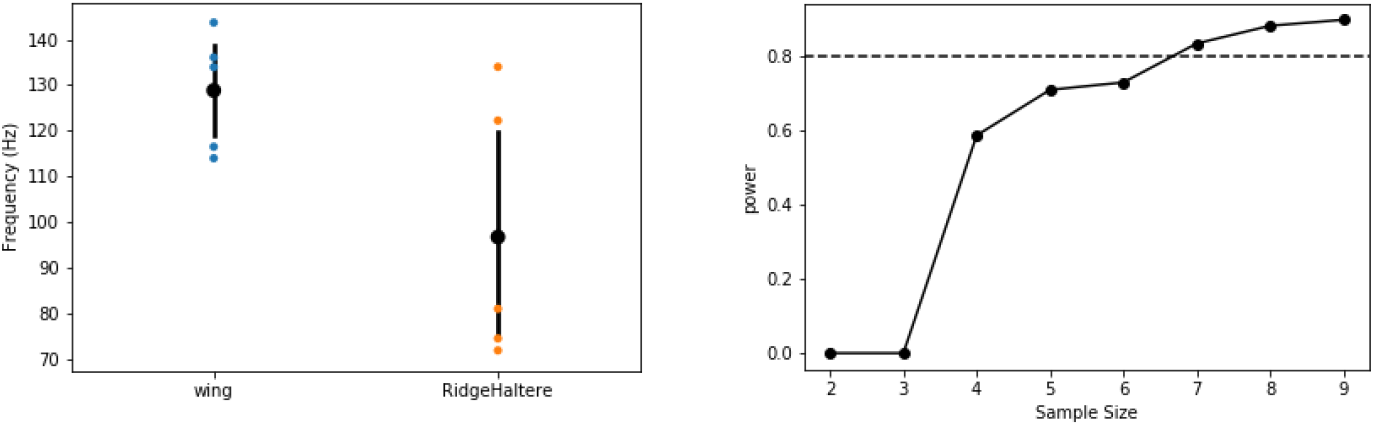

(*Left*) The frequency of wing and epimeral ridge cut haltere at wing length bin of (06 - 0.7) which has the largest difference) are significantly different (Wilcoxon signed-rank test, p = 0.039). (*Right*) Power analysis for different sample sizes. At our current sample size, we have a power of 0.713 that is we have a 71.3 % chance of picking up a significant difference.

### Integrity of Epimeral Ridge and Thoracic Resonance (Figure 4)

For one-way Kruskal Wallis test, our sample size, n = 8 for control and 6 for the 3 treatments groups each. We used a similar bootstrapping method, simulating our data based on our group mean and std, and calculating the power at difference sample sizes. To detect a significant difference at 80% chance we require a minimum of 9 samples. With our current sample size (n = 6), we have a substantial Type II error: that is a 48% chance of not detecting a difference if they were indeed different.

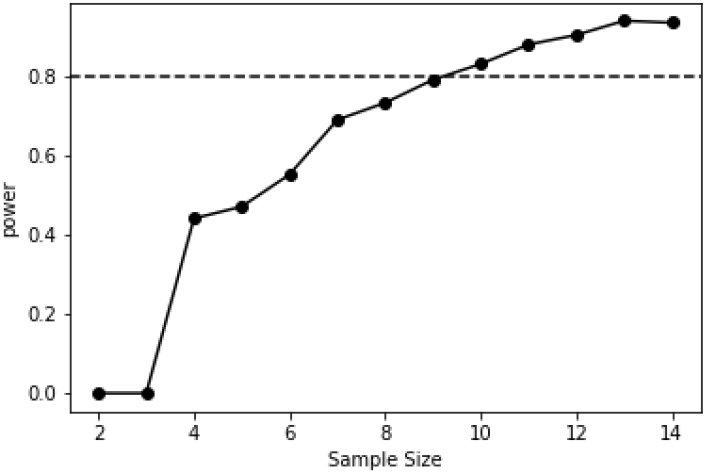

### Loading Haltere (Figure 5)

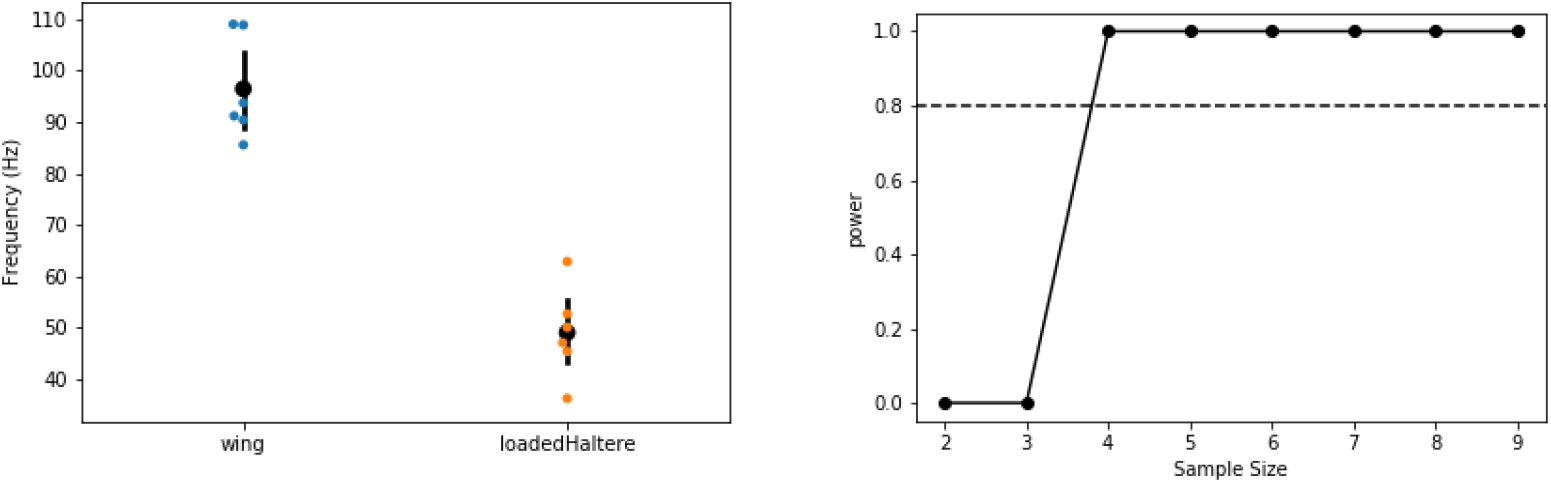

(*Left*) The frequency of wing and loaded haltere at maximum loading (“load3”) are significantly different (Wilcoxon signed-rank test (p =0.013). (*Right*) Power analysis for different sample sizes. Our sample size of 6 is greater than the minimum sample size (=4) needed to have 80% confidence level (dashed line)

---

**Supplementary Figure 1:**
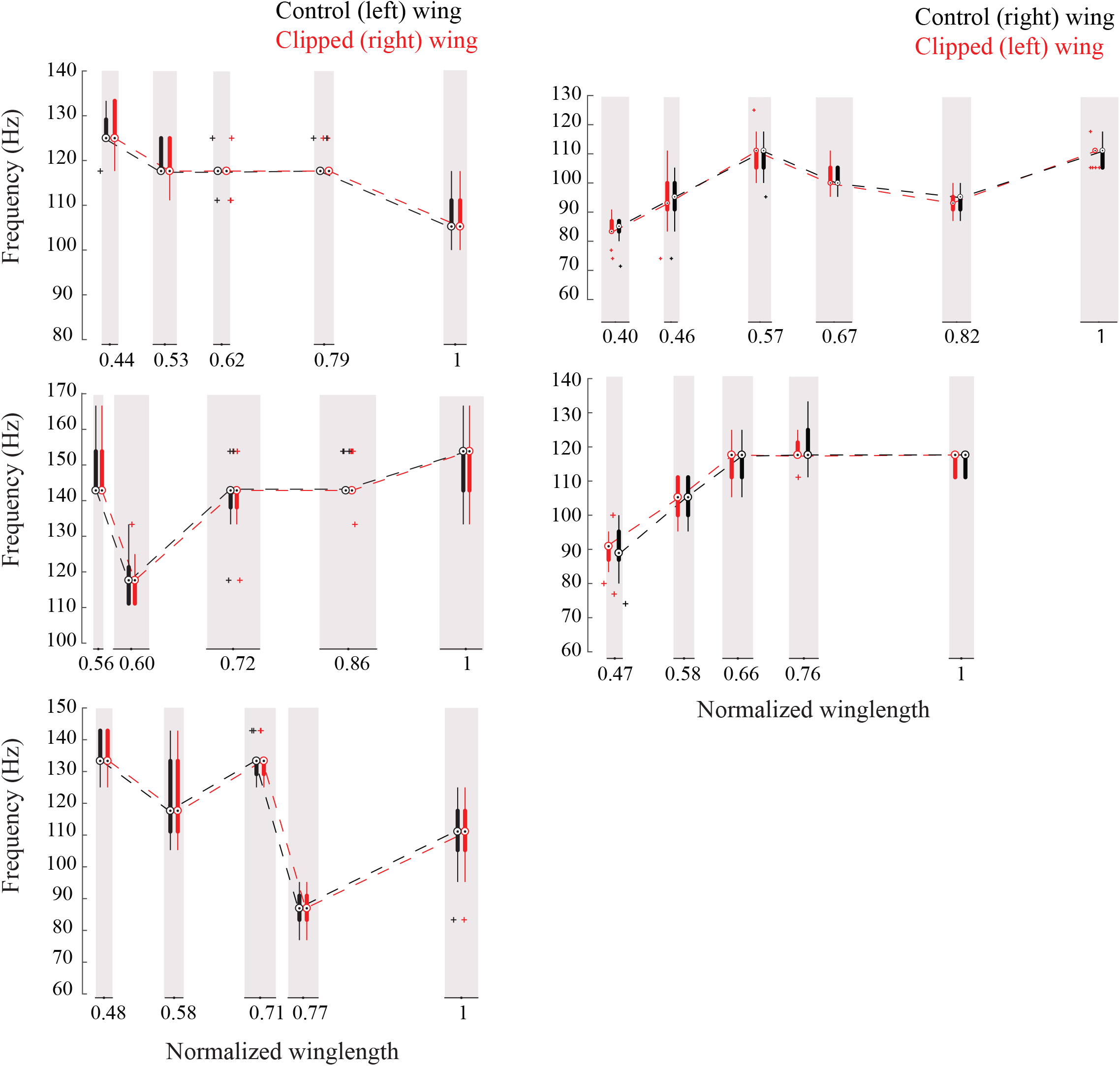
The frequencies of the two wings are coupled. Frequency of the intact (black) and clipped (red) wing plotted as compact box plot as a function of the clipped wing length for five individual flies showing that flies always flap the wings at the same frequency.

**Supplementary Figure 2:**
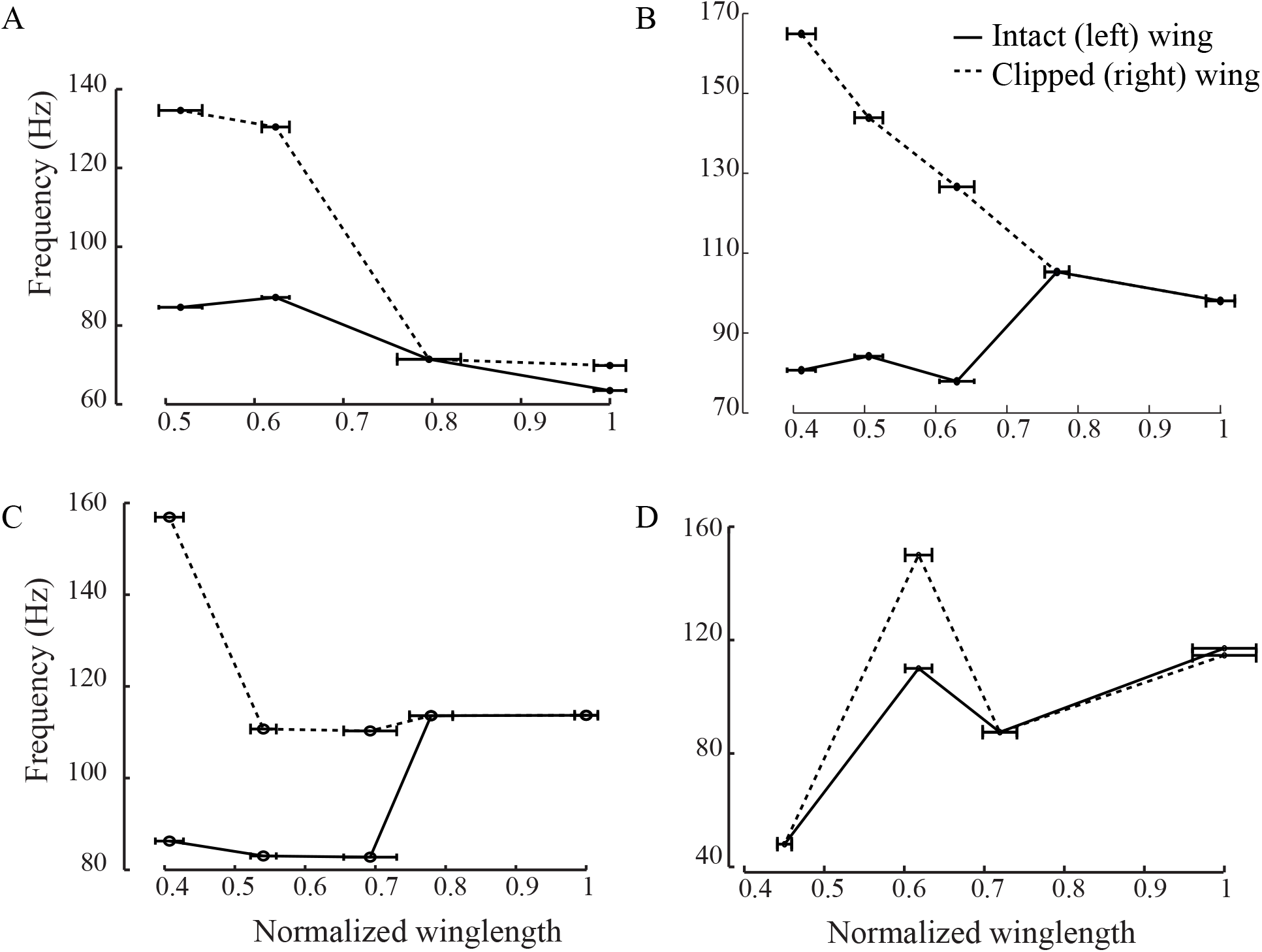
Scutellum synchronizes the frequencies of the two wings. Individual plots show the peak wing beat frequency for intact (black solid) and clipped (black dotted) wing as a function of the clipped wing length for four individual flies showing that the scutellum-lesioned flies flap their wings at different frequencies.

**Supplementary Figure 3:**
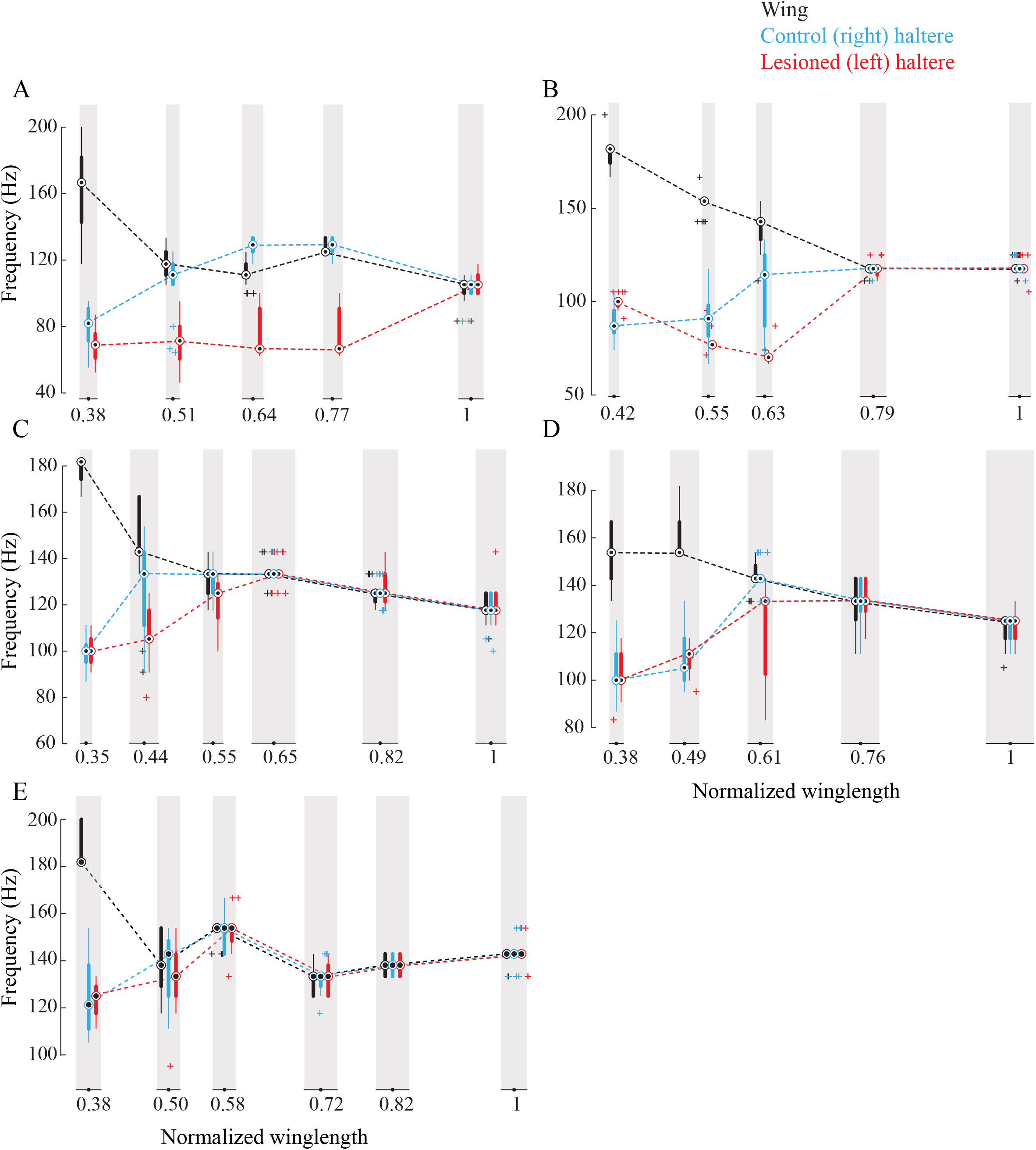
Sub-epimeral ridge couples the frequency of wings and halteres. (A-E) Frequency of wing (black), control haltere (blue) and haltere with the sub-epimeral ridge lesioned (red) plotted as compact box plot as a function of wing length for five individual flies.

**Supplementary Figure 4:**
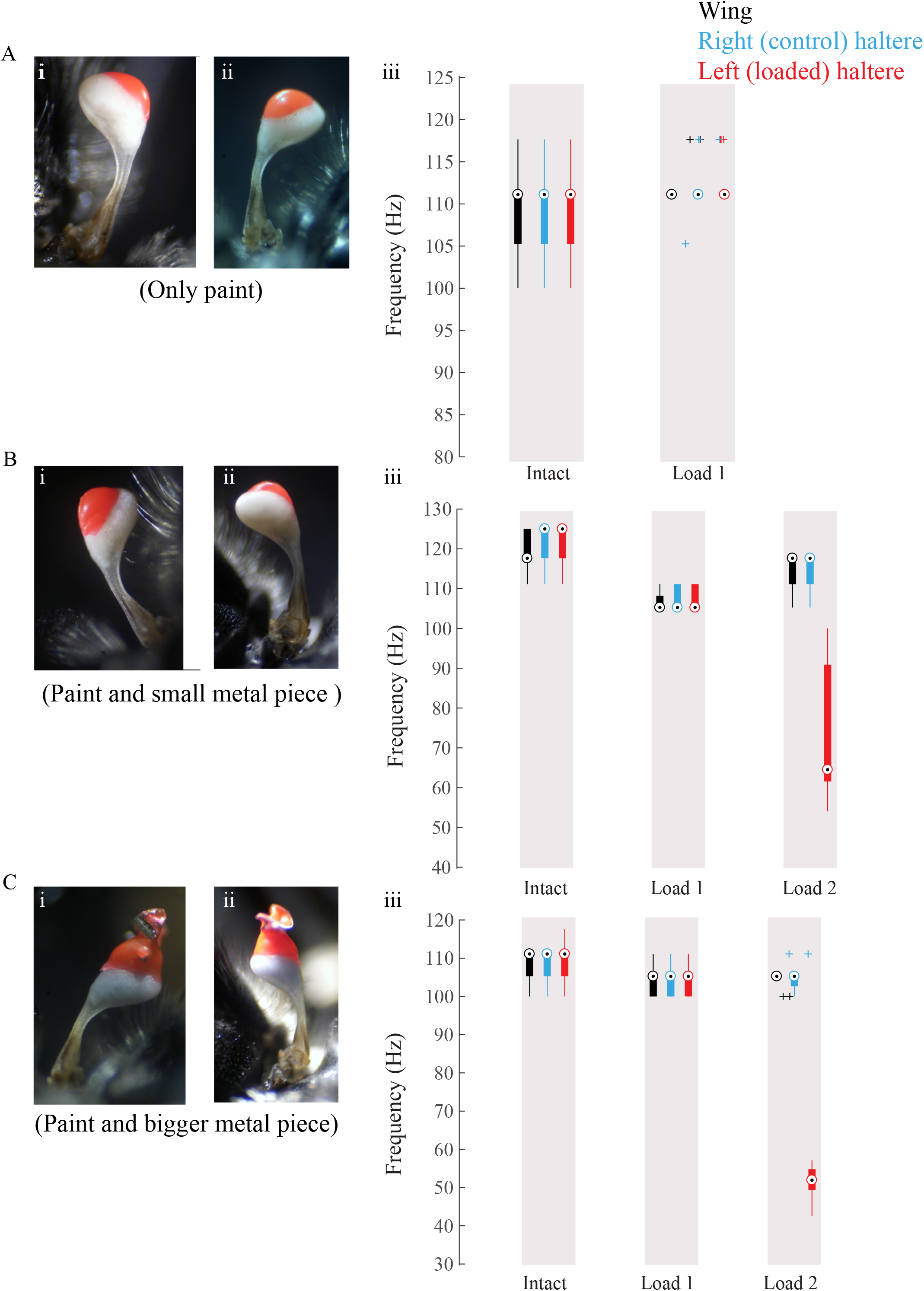
Increasing amounts of glue, paint and metal shavings were added to load halteres. (A-C) Data for individual flies showing loaded haltere (load visible in red) in the dorsal (i) and ventral (ii) views and the corresponding wing (black), control right haltere (blue) and loaded left haltere (red) frequency.

**Supplementary Figure 5:**
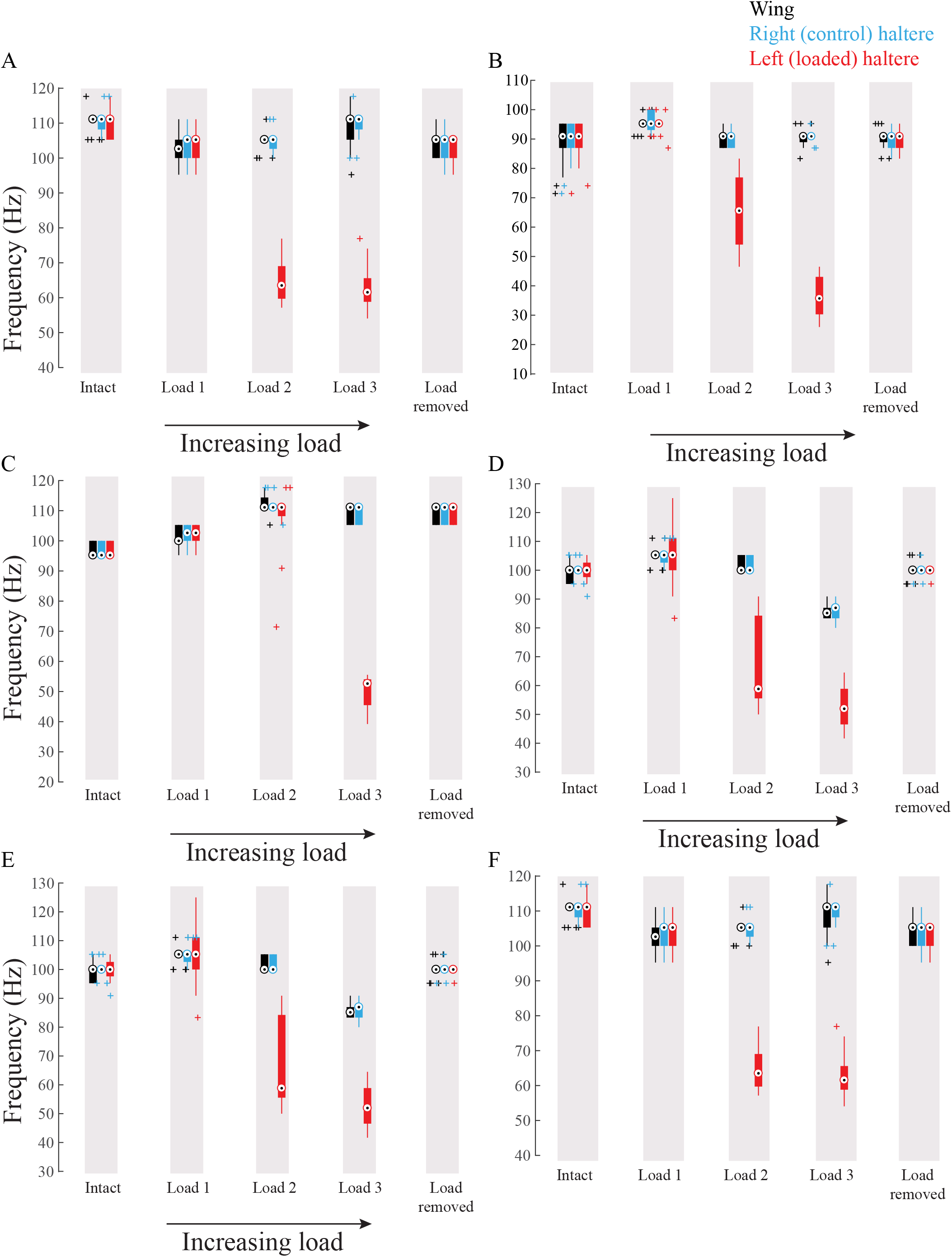
Wing haltere coupling acts unidirectionally such that changing haltere frequency does not influence wing frequency. (A-F) Frequency of wing (black), control haltere (blue) and loaded haltere (red) plotted as compact box plot for five individual flies. The haltere frequency drops as the haltere is loaded but the wing the frequency remains the same showing that wings and halteres coupling works in a unidirectional manner.

**Supplementary Figure 6:**
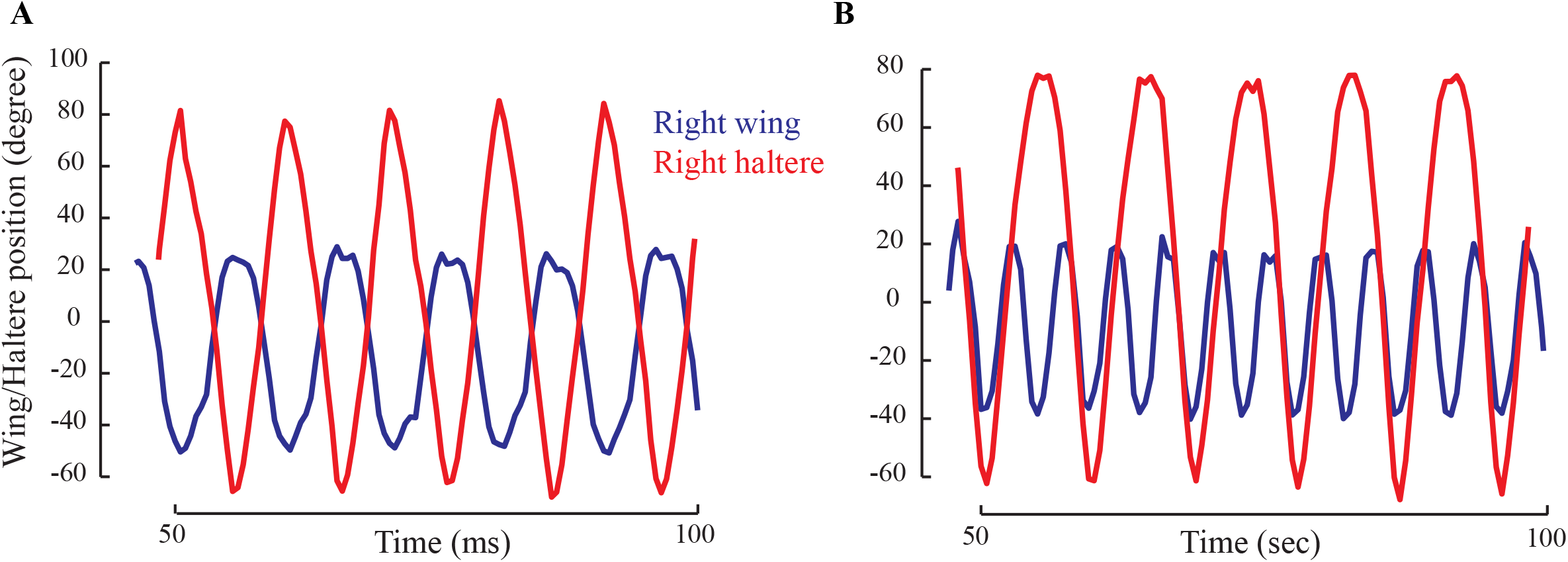
The positions of wing (blue) and haltere (red) showing their relative phase for a representative fly with intact wings (A) and wings cut (B). (A) When the wing and haltere frequencies are matched, they are perfectly antiphase. (B) However, when the beat frequency increases, the haltere cycles through different phases relative to the wing.

**Supplementary Figure 7:**
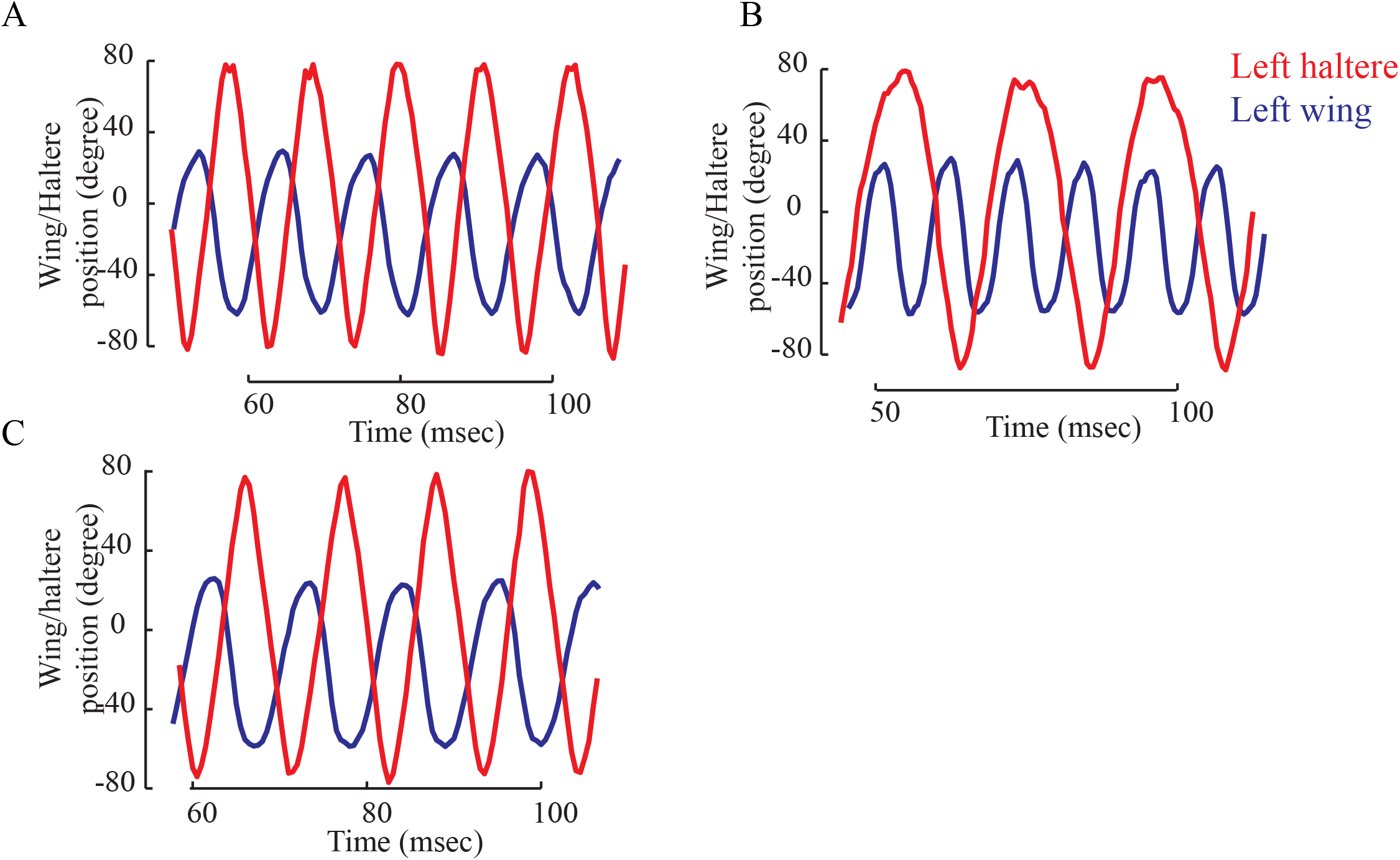
The wing (blue) and haltere (red) position showing their relative phase for a representative fly with intact halteres (A), loaded halteres (B) and the load removed (C). (A) The wing and haltere oscillate antiphase when their frequencies matches (A and C), but haltere cycles through different phases relative to the wing when the haltere frequency drops.

## References

Bender JA, Dickinson MH. 2006. Comparison of visual and haltere-mediated feedback in the control of body saccades in Drosophila melanogaster. J Exp Biol 209:4597–4606. doi:10.1242/jeb.02583

Cartar R V. 1992. Morphological Senescence and Longevity - an Experiment Relating Wing Wear and Life-Span in Foraging Wild Bumble Bees. J Anim Ecol 61:225–231. doi:10.2307/5525

Chan WP, Dickinson MH. 1996. Position-specific central projections of mechanosensory neurons on the haltere of the blow fly, Calliphora vicina. J Comp Neurol 369:405–418. doi:10.1002/(sici)1096-9861(19960603)369:3<405::aid-cne6>3.0.co;2-9

Combes S, Crall J, Mukherjee S. 2010. Dynamics of animal movement in an ecological context: dragonfly wing damage reduces flight performance and predation success. Biol Lett 6:426–429. doi:10.1098/rsbl.2009.0915

Deora T, Gundiah N, Sane SP. 2017. Mechanics of the thorax in flies. J Exp Biol 220. doi:10.1242/jeb.128363

Deora T, Singh AKAK, Sane SPSPS. 2015. Biomechanical basis of wing and haltere coordination in flies. Proc Natl Acad Sci 112:201412279. doi:10.1073/pnas.1412279112

Dickinson MH, Tu MS. 1997. The function of dipteran flight muscle. Comp Biochem Physiol a-Physiology 116:223–238. doi:10.1016/s0300-9629(96)00162-4

Dudley R. 2000. The Biomechanics of Insect Flight. New Jersey: Princeton University Press.

Ennos AR. 1987. A Comparative-Study of the Flight Mechanism of Diptera. J Exp Biol 127:355–372.

Esch H, Franz G. 1991. Neural Control of Fibrillar Muscles in Bees During Shivering and Flight. J Exp Biol 159:419–431.

Fayyazuddin A, Dickinson MH. 1999. Convergent mechanosensory input structures the firing phase of a steering motor neuron in the blowfly, Calliphora. J Neurophysiol 82:1916–1926.

Fayyazuddin A, Dickinson MH. 1996. Haltere afferents provide direct, electrotonic input to a steering motor neuron in the blowfly, Calliphora. J Neurosci 16:5225–5232.

Fernández MJ, Springthorpe D, Hedrick TL. 2012. Neuromuscular and biomechanical compensation for wing asymmetry in insect hovering flight. J Exp Biol 3631–3638. doi:10.1242/jeb.073627

Fox JL, Fairhall AL, Daniel TL. 2010. Encoding properties of haltere neurons enable motion feature detection in a biological gyroscope. Proc Natl Acad Sci U S A 107:3840–3845. doi:10.1073/pnas.0912548107

Fry SN, Sayaman R, Dickinson MH. 2003. The Aerodynamics of Free-Flight Maneuvers in Drosophila. Science (80-) 300:495–498.

Gordon S, Dickinson MH. 2006. Role of calcium in the regulation of mechanical power in insect flight. Proc Natl Acad Sci 103:4311–4315.

Haas CA, Cartar R V. 2008. Robust flight performance of bumble bees with artificially induced wing wear. Can J Zool 86:668–675. doi:10.1139/Z08-034

Hall JM, McLoughlin DP, Kathman ND, Yarger AM, Mureli S, Fox JL. 2015. Kinematic diversity suggests expanded roles for fly halteres. Biol Lett 11:20150845. doi:10.1098/rsbl.2015.0845

Hayes EJ, Wall R. 1999. Age-grading adult insects: A review of techniques. Physiol Entomol 24:1–10. doi:10.1046/j.1365-3032.1999.00104.x

Hedenström A, Ellington CP, Wolf TJ. 2001. Wing wear, aerodynamics and flight energetics in bumblebees (Bombus terrestris): An experimental study. Funct Ecol 15:417–422. doi:10.1046/j.0269-8463.2001.00531.x

Heide G, Götz KG. 1996. Optomotor Control of Course and Altitude in Drosophila Melanogaster is Correlated with Distinct Activities of at Least Three Pairs of Flight Steering Msucles. J Exp Biol 1726:1711–1726.

Hengstenberg R. 1993. Multisensory control in insect oculomotor systems. Rev Oculomot Res 5:285–98.

Land M, Collett T. 1974. Chasing Behaviour of Houseflies (Fannia canicularis). J Comp Physiol 331–357.

Lehmann F, Skandalis DA, Berthé R. 2013. Calcium signalling indicates bilateral power balancing in the Drosophila flight muscle during manoeuvring flight. J R Soc Interface 10.

Leston BYD, Pringle JWS, White DC. 1965. Muscular Activity During Preparation For Flight in a Beetle. J Exp Biol 42:409–414.

Lindsay T, Sustar A, Dickinson M. 2017. The Function and Organization of the Motor System Controlling Flight Maneuvers in Flies. Curr Biol 27:345–358. doi:10.1016/j.cub.2016.12.018

Miyan JA, Ewing AW. 1985. How Diptera move their wings - A re-examination of the Wing Base Articulation and Muscle Systems Concerned with Flight. Philos Trans R Soc London Ser B-Biological Sci 311:271–302. doi:10.1098/rstb.1985.0154

Mountcastle AM, Combes SA. 2014. Biomechanical strategies for mitigating collision damage in insect wings: structural design versus embedded elastic materials. J Exp Biol 217:1108–1115. doi:10.1242/jeb.092916

Nalbach G. 1994. Extremely Nonorthogonal Axes in a Sense Organ for Rotation - Behavioral-Analysis of the Dipteran Haltere System. Neuroscience 61:149–163. doi:10.1016/0306-4522(94)90068-x

Nalbach G. 1993. The Halteres of the Blowfly Calliphora .1. Kinematics and Dynamics. J Comp Physiol a-Sensory Neural Behav Physiol 173:293–300. doi:10.1007/bf00212693

Nalbach G, Hengstenberg R. 1994. The halteres of the blowfly Calliphora II. Three-dimensional organization of compensatory reactions to real and simulated rotations. J Comp Physiol 175:695–708.

Polilov AA. 2015. Small Is Beautiful: Features of the Smallest Insects and Limits to Miniaturization. Annu Rev Entomol 60:103–121. doi:10.1146/annurev-ento-010814-020924

Polilov AA. 2012. The smallest insects evolve anucleate neurons. Arthropod Struct Dev 41:29–34. doi:10.1016/j.asd.2011.09.001

Pringle JWS. 1949. The Excitation And Contraction Of The Flight Muscles Of Insects. J Physiol 108:226–232.

Pringle JWS. 1948. The Gyroscopic Mechanism Of The Halteres Of Diptera. Philos Trans R Soc London Ser B-Biological Sci 233:347–384. doi:10.1098/rstb.1948.0007

Sane SP. 2016. Neurobiology and biomechanics of flight in miniature insects. Curr Opin Neurobiol 41:158–166.

Sherman A, Dickinson MH. 2003. A comparison of visual and haltere-mediated equilibrium reflexes in the fruit fly Drosophila melanogaster. J Exp Biol 206:295–302. doi:10.1242/jeb.00075

Strogatz SH. 1994. Non-linear Dynamics and Chaos. New York: Perseus Books.

Trimarchi JR, Schneiderman AM. 1995. Different neural pathways coordinate Drosophila flight initiations evoked by visual and olfactory stimuli. J Exp Biol 198:1099–1104.

Vance JT, Roberts SP. 2014. The effects of artificial wing wear on the flight capacity of the honey bee Apis mellifera. J Insect Physiol 65:27–36. doi:10.1016/j.jinsphys.2014.04.003

Yarger AM, Fox JL. 2018. Single mechanosensory neurons encode lateral displacements using precise spike timing and thresholds. Proc R Soc B Biol Sci 285. doi:10.1098/rspb.2018.1759

## References

Hedrick TL. 2008 Software techniques for two- and three-dimensional kinematic measurements of biological and biomimetic systems. Bioinspir. Biomim. 3. (doi:10.1088/1748-3182/3/3/034001)

